# MaSkel and napari-MaSkel: fast morphological skeleton feature extraction from segmentation masks

**DOI:** 10.64898/2026.09.15.751651

**Authors:** Simon Wittmann, Dominik Pysch, Stefan Uderhardt, David B. Blumenthal, Anna Möller

## Abstract

Summary: Existing workflows for extracting morphological features from network-like biomedical structures often require several heterogeneous tools, which can limit integration and reproducibility. We present MaSkel and napari-MaSkel, open-source Python tools that provide GUI- and CLI-based workflows for 2D and 3D skeletonization and graph-based feature extraction, using a substantially accelerated implementation of the widely used Lee94 thinning algorithm. Availability and Implementation: MaSkel and napari-MaSkel are open-source Python packages available under the MIT license on PyPI and GitHub: https://github.com/bionetslab/maskel/, https://github.com/bionetslab/napari-maskel/. Documentation: https://bionetslab.github.io/maskel/, https://bionetslab.github.io/napari-maskel/. Benchmarking and validation: https://github.com/bionetslab/maskel-evaluations.

## 1. Introduction

Skeleton-based analysis is a common approach to quantify the geometry and topology of branching biological structures, such as vasculature, fibers, neurites, and other network-like objects. A typical workflow consists of three steps: thinning of binary masks into one-pixel-wide skeletons, interpreting the skeleton as a pixel graph, and extracting object-, branch-, or node-level morphological characteristics. These interpretable features enable diverse downstream analyses. For example, skeleton-derived centerlines, bifurcation points, and radii have been used to quantify whole-mouse 3D brain vasculature and reveal strain-specific differences in vascular density and branching [Todorov et al., 2020]. In 2D retinal images, skeleton-based graph descriptors of vessel geometry and connectivity have enabled disease classification [Budai et al., 2013]. In multi-label instance segmentations of cells (as in Schnitzerlein et al. [2025]), skeleton-based shape descriptors can be used to quantify protrusiveness and sampling dynamics to distinguish functional states (all three shown as examples in Figure 1).

**Figure 1.**
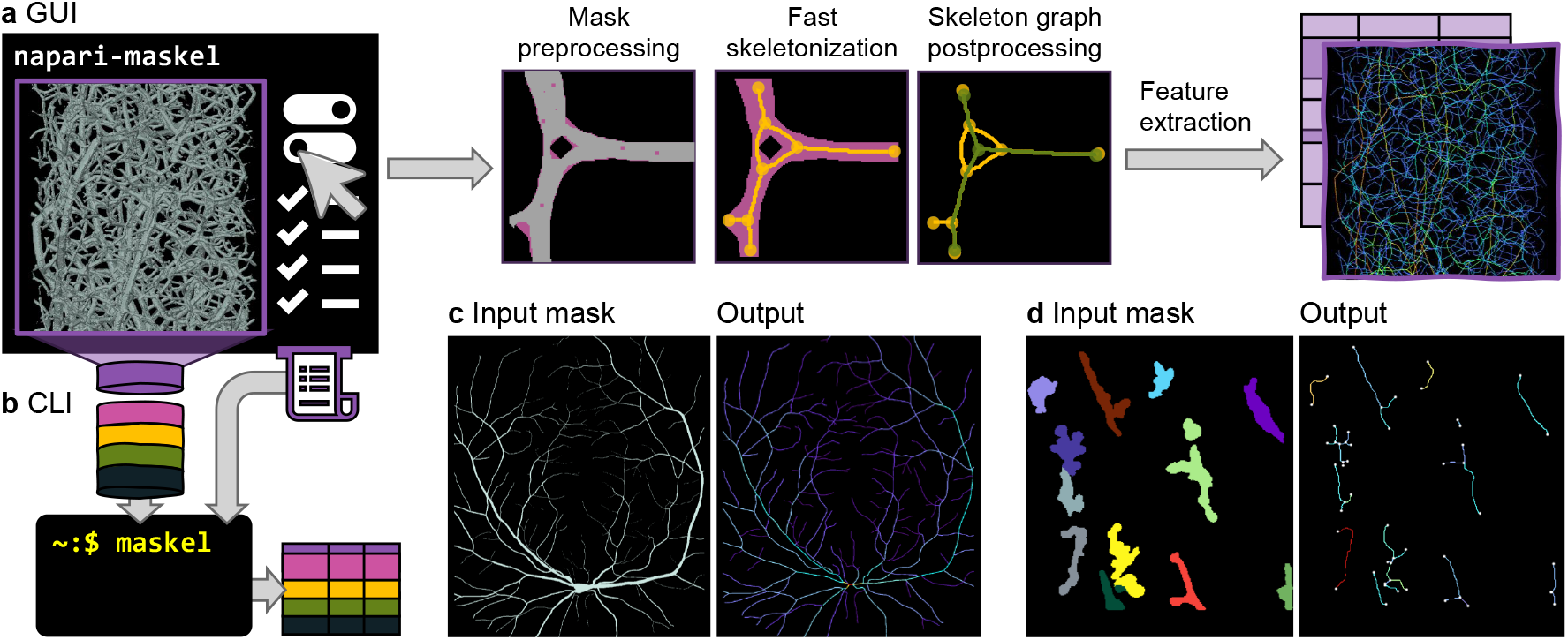
Interplay of napari-MaSkel and MaSkel. (a) napari-MaSkel allows to interactively configure the parameters for mask preprocessing, skeletonization, and postprocessing by visual inspection of intermediate steps and final output in the graphical user interface (GUI). (b) The resulting recipe can then be applied to an entire input batch with the command line interface (CLI) with MaSkel. (c) Application example 1: Retinal vessel mask [Budai et al., 2013] and output skeleton branches, colored by mean radius of each branch. (d) Application example 2: Instance-segmented macrophages [Schnitzerlein et al., 2025] and output skeleton keypoints and branches, colored by tortuosity.

Several software tools address different parts of this workflow, but each has limitations: The MATLAB-based tool REAVER [Corliss et al., 2020] and the FIJI macro TWOMBLI [Wershof et al., 2021] support only 2D data. PoreSpy [Gostick et al., 2019] targets porous materials rather than biomedical imaging, whereas Skan [Nunez-Iglesias et al., 2018] provides extensive graph-based feature extraction but requires a precomputed skeleton as input. VesSAP [Todorov et al., 2020] lacks a graphical user interface and is primarily targeted towards light-sheet fluorescence microscopy of vasculature. VIPAR [Hägerling et al., 2017] and VesselVio [Bumgarner and Nelson, 2022] are distributed as standalone applications and are therefore less easily integrated into Python-based image analysis workflows.

The napari ecosystem [Chiu et al., 2022] also contains plugins for skeletonization and skeleton-based feature extraction: napari-vessel-express [Spangenberg et al., 2023] focuses on segmentation parameter configuration that must then be run within a separate Docker image to retrieve feature output; napari-toska [Ryan et al., 2026] conceptually supports 3D graph representations but its graphical user interface (GUI) fails to process 3D volumes out-of-the-box due to dimensionality handling errors. An integrated workflow for skeleton generation, visualization, and feature extraction within napari is lacking.

Most Python-based workflows rely on the skeletonization routines provided by scikit-image [van der Walt et al., 2014], which implement the thinning algorithms of Zhang and Suen [1984] for 2D images and Lee et al. [1994] for 3D volumes (we refer to these algorithms as “Zhang84” and “Lee94”). These implementations are widely adopted but their runtime can become a limiting factor for large datasets. VesselVio [Bumgarner and Nelson, 2022] addresses this with a Numba-parallelized [Lam et al., 2015] implementation of Lee94 for 3D data but provides this speed-up only for 3D and not for 2D data.

To address these limitations, we present MaSkel and napari-MaSkel. MaSkel implements the algorithms for skeletonization and graph-based feature extraction and provides a command line interface (CLI); napari-MaSkel provides a GUI within the napari ecosystem. Both tools can independently execute the entire workflow from segmentation mask to feature table. Their combination enables users to interactively configure and evaluate a workflow on individual examples or smaller crops in the napari GUI and to then apply this workflow reproducibly to entire datasets via the CLI (Figure 1). For 3D data, MaSkel can compute over 70 features, spanning graph topology (e.g., node/edge counts, branch points), morphometry (length, tortuosity, radius, diameter, volume, surface area, fractal dimension), and spatial position (pixel/voxel coordinates). Under the hood, MaSkel implements a customized version of Lee94, which we accelerate by implementing provably equivalent but substantially faster custom subroutines for 2D and 3D data. By integrating skeleton generation, visualization, and feature-table output in a single workflow, napari-MaSkel and MaSkel reduce manual processing steps and facilitate reproducible skeleton-based quantification across diverse image modalities. The interoperability of napari further provides access to various tools for generating the segmentation masks required as input.

## 2. MaSkel and napari-MaSkel

For both tools, the workflow is configured through a recipe, which can be edited directly as json file or interactively in napari-MaSkel. The recipe specifies the physical spacing of the input image, including anisotropic pixel or voxel sizes, and the desired feature extraction layers. Object-level, branch, and node features can be selected independently, and branch labels and color-coding by individual features can be displayed in the napari viewer. Several optional cleanup steps are available, including filling holes up to a specified maximum size, morphological closing, removal of small skeleton junctions, and iterative pruning of short skeleton spurs (Appendix A). The calculation of computationally more expensive fractal-dimension and radius-related features can be enabled separately. Output settings control how the skeleton and the extracted features are saved.

A binary mask, such as a semantic segmentation of vasculature or neurites, is processed as a single network. Multi-label masks, such as cell instance segmentations, are instead processed independently for each object ID. Hence, objects (pixels or voxels sharing the same ID) are skeletonized separately, to preserve individual shape characteristics, rather than skeletonizing the entire foreground and then assigning the IDs (as done e.g. in Ryan et al. [2026]). The original object IDs are retained in the resulting feature tables.

We generally recommend using napari-MaSkel to configure the recipe. When working with a single sample, this provides full control over the intermediate steps of the workflow. Overlaying the original signal, mask, and skeleton allows biomedical experts to visually assess whether the resulting connections and shapes are biologically coherent. For large imaging volumes, a smaller crop can instead be used to efficiently inspect the effects of preprocessing, skeletonization, and graph cleanup. When processing an entire dataset containing multiple masks, a representative sample can be selected to configure and evaluate the recipe before applying the finalized recipe to the complete dataset. This workflow can be summarized as follows:

1. The 2D or 3D segmentation mask is loaded into napari, which natively supports a wide range of image formats or provides plugins for less common formats. Numerous plugins also exist for binary or multi-label segmentation, in case the user starts with an intensity image. The mask must be converted to a labels layer.
2. Once installed, napari-MaSkel can be opened from napari’s “Plugins” menu by clicking “Analyze mask (MaSkel)”.
3. Load an existing recipe or configure the desired parameters, and save the recipe for later use.
4. Click “Analyze mask” to execute the workflow and display the selected intermediate results and outputs in napari.
5. Inspect the skeleton and feature tables and iteratively refine the recipe if required. Object-level, branch, and node features can be inspected using napari’s “Feature table widget” from the “Plugins” menu.
6. Apply the finalized recipe to additional masks with MaSkel from the command line by specifying the input directory, output directory, and recipe.

For a qualitative validation, we used the HRF dataset [Budai et al., 2013] which comprises 45 manually generated vessel masks of retinal fundus images balanced across healthy individuals, diabetic retinopathy patients, and glaucoma patients. We trained several machine learning models to predict disease states from MaSkel’s sample-level skeleton features. In a 5-fold cross-validation, logistic regression achieved the highest accuracy (86.7%) (Appendix A).

## 3. Accelerating Lee94

Lee94 was originally formulated for 3D volumes: it examines the 3 × 3 × 3 neighborhood of each foreground voxel and iteratively removes voxels in six sub-iterations (north, south, east, west, up, down) until convergence. A voxel is removed only if (i) it is not an endpoint (exactly one foreground neighbor, marking a branch tip), (ii) its removal leaves the Euler characteristic invariant (no holes or tunnels created or destroyed), and (iii) it is a simple point (number of connected foreground components is preserved). Because removing one voxel can affect the simplicity of its neighbors, each sub-iteration first collects all admissible candidates and then removes them sequentially, rechecking candidates whose neighborhood has changed.

Most Python-based tools build on scikit-image’s `skeletonize`, which defaults to Zhang84 for 2D and to Lee94 for 3D data. Lee94 can also be used for 2D input, in which case the image is padded into a single-slice 3D volume and processed by the 3D routine. Although two z-direction sub-iterations are skipped, the Euler-invariant and simple-point checks still operate on the 26-neighborhood. The whole routine is single-threaded. VesselVio [Bumgarner and Nelson, 2022] also relies on scikit-image for 2D but provides a custom Numba-accelerated [Lam et al., 2015] Lee94 implementation for 3D that parallelizes candidate identification and removability evaluation.

MaSkel implements dedicated 2D and 3D routines that exploit dimensionality-specific properties of Lee94. For 2D, it uses a 4-neighbor border check and an 8-neighbor simple-point test, omitting the redundant Euler-invariance check. The simple-point test is implemented as a lookup into a precomputed table of all 2^8^ = 256 8-neighbor configurations. Candidate identification and removability evaluation are parallelized with Numba, removal remains sequential. These simplifications preserve the removal decisions of Lee94, so the 2D output is bit-identical (proofs in Appendix B).

For 3D data, MaSkel retains the full test. Candidate identification is parallelized per z-slice via a two-pass counting and prefix-sum scheme, removability evaluation is parallelized across candidates. During removal, a candidate’s simplicity is rechecked only if one of its 26 neighbors was removed earlier in the same pass, avoiding redundant checks. The simple-point test uses a precomputed 26-neighbor adjacency structure and an explicit stack for connected component labeling.

We benchmarked our accelerated Lee94 implementation on two public datasets: the 2D HRF dataset described above and the VESSEL12 dataset [Rudyanto et al., 2014] which contains 20 thoracic CT scans with ground-truth lung masks from a segmentation challenge. As competitors, we included scikit-image’s Zhang84 implementation (only for 2D data), scikit-image’s Lee94 implementation (for both 2D and 3D data), and VesselVio’s Lee94 implementation (only for 3D data).

On HRF, MaSkel was on average 11.83 times faster than scikit-image-Lee94 and 3.40 times faster than scikit-image-Zhang84 (Table 1). On VESSEL12, it was 3.17 and 4.14 times faster than scikit-image-Lee94 and VesselVio-Lee94, respectively. Note that VesselVio’s parallelized Lee94 implementation was thus slower than scikit-image’s sequential version, possibly due to suboptimal usage of recursion in VesselVio. MaSkel’s increased parallelism incurred a modest memory overhead relative to scikit-image, but it still used less memory than VesselVio (Appendix A). Skeletons from MaSkel’s and scikit-image’s Lee94 implementations were bit-identical (Appendix A), in line with our theoretical considerations.

**Table 1.** Runtimes (mean ± std across samples) of MaSkel’s Lee94 implementation, scikit-image’s Zhang84 and Lee94 implementations, and VesselVio’s Lee94 implementations. The implementations were tested on 45 2D images from the HRF dataset (2336 × 3504 pixels with 631 094 ± 115 946 foreground px per image) and 20 3D scans from the VESSEL12 dataset ((430 ± 48) × 512 × 512 voxels with 13 525 543 ± 4 424 202 foreground voxels per scan).

| Dataset | Method | Runtime (s) |
| --- | --- | --- |
| HRF (2D) | MaSkel (Lee94) | **$0.09 \pm 0.01$** |
| | scikit-image (Zhang84) | $0.31 \pm 0.06$ |
| | scikit-image (Lee94) | $1.09 \pm 0.23$ |
| VESSEL12 (3D) | MaSkel (Lee94) | **$9.76 \pm 2.77$** |
| | scikit-image (Lee94) | $30.40 \pm 6.77$ |
| | VesselVio (Lee94) | $41.20 \pm 15.13$ |

## 4. Discussion

We present MaSkel and napari-MaSkel, a Python-based workflow for skeletonization and quantitative analysis of branching biological structures. Both tools can independently execute the complete workflow from binary or multi-label masks to object-, branch-, and node-level features. Their combination enables users to configure and inspect a workflow interactively in napari and subsequently apply the same recipe reproducibly to entire datasets. The custom implementation of Lee94 thinning produces bit-identical skeletons to established implementations while substantially reducing runtime for both 2D and 3D data.

An advantage of MaSkel is its flexibility regarding biological structures and imaging modalities. Vessel segmentation masks can be used to quantify vascular morphology, including branch lengths, junction properties, and network-level characteristics. Neurite segmentations can be analyzed to quantify branching and process morphology, while fiber-like structures can be characterized by their length, orientation, branching, and connectivity. Multi-label masks further allow individual objects to be skeletonized and quantified independently. The combination of intermediate visualization and tabular feature extraction enables domain experts to assess the resulting skeleton before applying a finalized recipe to larger datasets.

MaSkel is intended for the quantitative analysis of binary or multi-label segmentation masks and does not replace the segmentation step itself. The quality of the resulting skeleton and extracted features therefore depends on the quality of the input mask. Preprocessing and graph cleanup parameters can affect both topology and morphology and need to be selected with respect to the biological structures under investigation and the quality of the masks. While napari-MaSkel facilitates this process by allowing intermediate results to be inspected interactively, expert assessment remains necessary to determine whether the resulting skeleton is biologically meaningful.

Although our accelerated Lee94 implementation substantially improves runtime, very large 3D volumes can still require considerable computational resources. In future work, this limitation could be addressed through further optimizing our Lee94 implementation, e. g., by reducing repeated candidate scans and maintaining a persistent candidate set across sub-iterations. Moreover, a tiling-based approach may eventually be necessary for very large microscopy volumes. The modular design of MaSkel, together with napari’s extensibility, provides a basis for such extensions while retaining the same workflow.

## Acknowledgements

The authors gratefully acknowledge the scientific support and high-performance computing resources provided by the Erlangen National High Performance Computing Center of the Friedrich-Alexander-Universität Erlangen-Nürnberg. The hardware was funded by the German Research Foundation. Claude Code (Anthropic) was used to assist with code and manuscript review.

## Author contributions

S.W. (Methodology [lead], Software [lead], Validation [equal], Writing – original draft [equal]), D.P. (Software [supporting], Formal analysis [equal]), S.U. (Funding acquisition [equal], Supervision [equal]), D.B.B. (Funding acquisition [equal], Supervision [equal], Writing – review & editing [lead]), A.M. (Conceptualization [lead], Data curation [lead], Formal analysis [equal], Software [supporting], Supervision [equal], Validation [equal], Writing – original draft [equal]).

## Supplementary material

Supplementary material is available online. HRF and VESSEL12 are publicly available (see references).

