## Appendices for "MaSkel and napari-MaSkel: fast morphological skeleton feature extraction from segmentation masks"

Simon Wittmann[1], Dominik Pysch[1], Stefan Uderhardt[2,3,4], David B. Blumenthal[1], and Anna Möller[1,2,*]

[1] Biomedical Network Science Lab, Department Artificial Intelligence in Biomedical Engineering, Friedrich-Alexander-Universität Erlangen-Nürnberg, Erlangen, Germany
[2] Department of Medicine 3 – Rheumatology and Immunology, Friedrich-Alexander-Universität Erlangen-Nürnberg and Universitätsklinikum Erlangen, Erlangen, Germany
[3] Deutsches Zentrum für Immuntherapie, Friedrich-Alexander-Universität Erlangen-Nürnberg, Erlangen, Germany
[4] Exploratory Research Unit, Optical Imaging Competence Center Erlangen, Friedrich-Alexander-Universität Erlangen-Nürnberg, Erlangen, Germany

### A Supplementary results

#### A.1 Pipeline: preprocessing, cleanup, and feature extraction

*Preprocessing.* Segmentation masks can contain small holes and gaps, particularly at faint structures or vessel crossings; left untreated, the thinning algorithm traces circular paths around these holes' boundaries, producing spurious loops in the resulting skeleton graph. MaSkel therefore offers two optional preprocessing steps, applied before thinning: morphological closing (`scipy.ndimage.binary_closing`) and hole filling (`scipy.ndimage.binary_fill_holes`), which fills every enclosed background region in the mask. A configurable `max_hole_size` threshold reverts filled holes above a given area/volume, so that legitimate anatomical voids are preserved.

*Postprocessing.* Two optional cleanup steps operate on the skeleton after thinning. Junction cleanup targets small triangle/diamond artifacts at bifurcations, a well-known thinning side effect: the skeleton is converted to a graph via Skan [Nunez-Iglesias et al., 2018], every elementary cycle is enumerated (`networkx.cycle_basis`), and a cycle is collapsed to a single centroid pixel whenever its perimeter is smaller than the configurable parameter `cleanup_threshold_factor` (default 2.5) times the local diameter estimated from the Euclidean distance transform at the cycle's nodes — a real bifurcation has a wide perimeter relative to the local vessel diameter, while a spurious triangle artifact is small and tight. Overlapping small cycles are merged before collapsing, and external branches are reconnected to the new centroid via Bresenham line rasterization to preserve connectivity. Separately, spur pruning removes short branches that connect an endpoint (degree 1) directly to a junction (degree >1) and fall below a configurable `min_spur_length` — also thinning artifacts rather than true branch tips — repeating for a configurable number of iterations, since removing one spur can expose another beneath it.

*Feature extraction.* The (optionally cleaned) skeleton is converted into a graph via Skan's `Skeleton` object, which builds a sparse adjacency matrix over foreground pixels/voxels; `skan.summarize` then decomposes this graph into branches (edges between endpoint/junction nodes), giving each branch's path length (`branch-distance`) and straight-line endpoint distance (`euclidean-distance`), from which tortuosity and straightness follow directly. Node degrees, endpoint/bifurcation counts, and the number of connected components (via `scipy.sparse.csgraph.connected_components` on a graph reconstructed from the branch endpoint pairs) give the topological features. Beyond what Skan itself provides, MaSkel adds several custom features: mask area and mask area fraction (foreground count, and its fraction of the crop); the hyphal growth unit (total branch length divided by endpoint count); fractal dimension, estimated via box-counting at scales $2^k$ up to $2^{\lfloor \log_2(\min(\text{shape})/4) \rfloor}$, with the $R^2$ of the log-log fit reported alongside it; and radius/diameter statistics from the Euclidean distance transform, sampled either over the whole object or per branch, with each branch additionally modeled as a chain of frustums between consecutive skeleton pixels to estimate its volume and surface area. All length-, area-, and volume-derived features above are reported in physical units whenever a pixel/voxel spacing is supplied, since Skan's underlying graph is itself built with that spacing; dimensionless ratios (tortuosity, straightness, mask area fraction) are correctly left unaffected by spacing. Fractal dimension — a scale-invariant log-log slope — is instead forced to zero whenever spacing is anisotropic, since box-counting assumes isotropic voxels.

##### A.2 Phenotype classification on HRF data

Using MaSkel's skeleton-derived features (per-image summary statistics spanning topology, morphology, and vascular density) extracted from all 45 HRF images (15 healthy, 15 diabetic retinopathy, 15 glaucoma; from https://www5.cs.fau.de/research/data/fundus-images/), we evaluated their ability to discriminate the three phenotypes with five classifiers: a support vector machine (SVM), logistic regression, a random forest, a hard-voting ensemble, and a stacking ensemble (logistic regression as meta-learner). Zero-variance features were removed (fit on the full 45-sample dataset, since this step is unsupervised); the remainder was reduced to the top ten by one-way analysis of variance (ANOVA) F-statistic (`SelectKBest`) and standardized to zero mean and unit variance within each model's own pipeline, refit on each training fold only, before hyperparameter-tuning via grid search under stratified 5-fold cross-validation. Logistic regression achieved the highest mean cross-validated accuracy (86.7%), ahead of random forest, an RBF-kernel SVM, and both the voting and stacking ensembles (82.2% each, a four-way tie). Feature importance from the random forest broadly agreed with the ANOVA ranking on the strongest predictors — vessel diameter/radius, mean segment volume, mask area (fraction), and mean surface area ranked among the top contributors in both — but the two rankings disagreed on mean tortuosity, which random forest ranked third most important while ANOVA ranked it least important among the ten selected features.

##### A.3 Empirical verification of equivalence between MaSkel's and scikit-image's Lee94 implementations

To empirically verify that MaSkel's Lee94 implementation and the one provided by the scikit-image package (`skimage.morphology.skeletonize(image, method="lee")`) produce equivalent output, we compared their output skeletons via exact array equality, on every image or scan in the two datasets used for the runtime benchmark: the 45 HRF images (2D) and the 20 VESSEL12 scans (3D; from https://zenodo.org/records/8055066). All 45 HRF images and all 20 VESSEL12 scans were bit-identical to scikit-image's output, with zero mismatched pixels or voxels in every case.

Beyond these datasets, MaSkel's test suite continuously verifies equivalence against scikit-image: two regression tests compare full skeletons on real image data (a black-and-white silhouette image for 2D, a 10-slice volumetric test scan for 3D, both from scikit-image's example data). Further fourteen hand-constructed patterns (nine in 2D, five in 3D) target edge cases behind Lemma B.1 and Lemma B.2 (holes, cavities, branch junctions, diagonal connectivity configurations). All of these checks assert exact array equality and run as part of MaSkel's continuous integration.

##### A.4 Memory benchmark of Lee94 implementations

Peak memory usage of the different thinning implementations was measured as the maximum resident set size (RSS) observed while each method executed, sampled every 10 ms from `/proc/self/status` on a background thread; each sample–method pair was run in a dedicated subprocess to prevent memory retained from a previous run from inflating later measurements. Active RSS polling was used instead of the OS-reported per-process high-water mark (`ru_maxrss`), which on Linux is seeded from the parent process's RSS at fork time and would otherwise conflate the benchmarking harness's own memory footprint with that of the method being measured. Each benchmark ran with exclusive access to a dedicated node of a compute cluster (dual-socket AMD EPYC 7502 "Rome" CPUs, 64 cores / 128 threads, 512 GB RAM per node), so no other jobs could contend for CPU or memory during measurement.

Table A.1: Peak memory usage (mean ± std) of MaSkel compared to scikit-image and VesselVio's Lee94 thinning implementations.

| Dataset | Method | Peak RAM (MB) |
|---|---|---|
| HRF ($n = 45$) | MaSkel (Lee94) | 205.34 ± 10.44 |
| | scikit-image (Zhang84) | 162.52 ± 2.06 |
| | scikit-image (Lee94) | 171.98 ± 0.19 |
| VESSEL12 ($n = 20$) | MaSkel (Lee94) | 510.18 ± 35.14 |
| | scikit-image (Lee94) | 458.88 ± 36.30 |
| | VesselVio (Lee94) | 735.44 ± 121.89 |

### B Redundancy of the Euler invariant check in 2D Lee94 thinning

#### B.1 Background

The Lee94 thinning algorithm [Lee et al., 1994] preserves an object's topology by iterating over all border points - points that lie on the current surface of the object - and checking if they satisfy *all three* of the following conditions:

1. **Non-endpoint**: the point must not have exactly one foreground neighbor. A pixel with exactly one foreground neighbor is the tip of a thin branch; protecting it stops thinning from eating away whole branches instead of just thickness.
2. **Euler invariant**: removing the point must not change the Euler characteristic of the image — i.e. it must not create or destroy a hole (or, in 3D, a tunnel/cavity).
3. **Simple point**: removing the point must not disconnect the foreground object.

In 3D, conditions 2 and 3 are complementary and independently necessary: the Euler-invariant check catches topology changes to holes/tunnels/cavities, while the simple-point check catches disconnection of the object itself — neither implies the other in general 3D. The remainder of this appendix shows that for 2D images, this independence collapses: condition 2 becomes automatically satisfied whenever condition 3 holds, so it can be dropped entirely.

#### B.2 2D Lee94

In 3D, a point $x$ is a border point if at least one of its 6 face-neighbors is background. In 2D, the border-point test is restricted to $N_4(x) \subset N(x)$, the 4 face-touching in-plane neighbors of $x$ (N, S, E, W). The effective border-point test becomes:

$$\exists\, y \in N_4(x) : y \in S^c,$$

i.e. $x$ is treated as a border point exactly when at least one of its 4 neighbors is background. This is the only sense in which "border point" is used for the remainder of this appendix. Furthermore, while in 3D both holes (tunnels) and cavities (hollow voids) contribute differently to the Euler characteristic of an object, cavities do not exist in 2D because an enclosed void and a through-hole are topologically identical.

In the following, let $O$ denote the total number of foreground connected components in the whole image. The background $S^c$ always splits into exactly one *unbounded* component (the "exterior", extending outward from the image) plus zero or more *bounded* components; a *hole* is one such bounded background component, and $H$ denotes their total count. The (2D) Euler characteristic is $\chi = O - H$ — one combined number summarizing how many separate pieces the object has and how many holes perforate it. We write $\Delta O$, $\Delta H$, $\Delta\chi$ for the change in $O$, $H$, $\chi$, respectively, caused by removing a single pixel $x$ from $S$.

Counting these components requires fixing how "connected" is defined for foreground versus background: throughout, foreground connectivity uses 8-adjacency (two foreground pixels are in the same component if reachable through a chain of 8-neighbors) while background connectivity uses 4-adjacency (only face-adjacent steps). This asymmetric $(8, 4)$ pairing is the standard convention in digital topology, and is necessary to avoid a paradox: if both used 8-connectivity, a single diagonal line of foreground pixels could simultaneously "cut" and "fail to cut" a diagonal line of background pixels crossing it at the same corner.

##### B.3 Impossibility of hole creation

**Lemma B.1 (Impossibility of hole creation).** *In 2D Lee94 thinning, removing a border point can never increase the number of holes ($\Delta H > 0$ is impossible).*

*Proof.* Suppose $x$ is removed (so its pixel becomes background). For this to *create* a new hole, the newly-background pixel at $x$'s location would have to become its own isolated bounded background component — meaning it must not be 4-adjacent to any pixel that was already background before the removal.

*Case 1: some $y \in N_4(x)$ is already background.* Then, after removing $x$, the pixel at $x$ is 4-adjacent to $y$, so it immediately joins $y$'s existing background component (whether that component is the exterior or an already-existing hole) rather than forming an isolated new one. No new hole is created:

$$\exists\, y \in N_4(x) : y \in S^c \implies \text{no hole created.}$$

*Case 2: every $y \in N_4(x)$ is foreground*, i.e. $\forall\, y \in N_4(x) : y \in S$. This is exactly the condition under which $x$ *fails* to be a border point — so $x$ is never even considered as a deletion candidate, and this case cannot occur for a pixel that Lee94 actually removes.

Since removal only ever happens in Case 1, no border-point removal can create a hole.

##### B.4 Impossibility of hole elimination

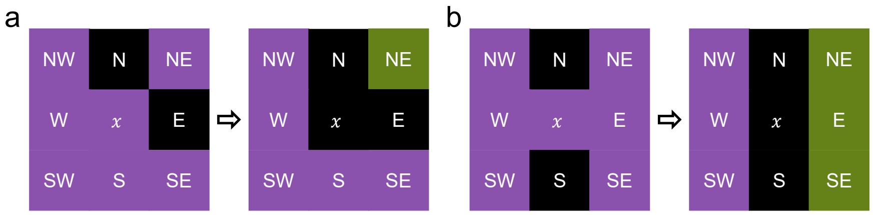


Fig. B.1: In 2D, the removal of a simple point cannot merge (a) two adjacent holes or (b) two opposite holes without increasing the number of connected components (marked green and purple).

**Lemma B.2 (Impossibility of hole elimination).** *In 2D Lee94 thinning, removing a simple point can never decrease the number of holes ($\Delta H < 0$ is impossible).*

*Proof.* A hole count can only decrease if removing $x$ *merges* two background components that were previously disconnected from each other, call them $B_1$ and $B_2$. This covers both ways a hole can disappear: a hole's interior merging with the exterior, or two distinct holes merging with each other. In either case, $x$ is, by assumption, the sole local separator between $B_1$ and $B_2$ — meaning $B_1$ and $B_2$ touch $x$'s neighborhood but are not otherwise connected to each other.

Since $B_1$ and $B_2$ only become connected once $x$ itself turns to background, they must occupy two of $x$'s 4-neighbors $N_4(x) = \{N, S, E, W\}$ — these are the only pixels that become newly 4-adjacent to $x$ after its removal (the diagonal/corner neighbors are never 4-adjacent to $x$, so they play no role in creating this particular connection).

We use one more geometric fact about the 3×3 neighborhood, concerning adjacency *among the neighbor cells themselves* (not their adjacency to $x$, which holds trivially for all of them by definition of $N(x)$): each corner neighbor $NE$, $SE$, $SW$, $NW$ is 8-adjacent to exactly two *other* cells of $N(x)$, namely its two flanking face-neighbors. There are exactly two ways two of the four face-neighbors can be arranged relative to each other, and we check both:

- *$B_1$, $B_2$ occupy adjacent face-neighbors* (e.g. $B_1$ at $N$, $B_2$ at $E$ in Figure B.1a). The corner cell $NE$ that flanks both of them cannot itself be background: if it were, it would already 4-connect $N$ and $E$ (and hence $B_1$ and $B_2$) directly, without needing $x$ removed at all — contradicting that $x$ is their sole

separator. So $NE$ must be foreground. But its only two possible neighbors in $N(x)$, namely $N$ and $E$, both belong to $B_1 \cup B_2$ (background). Hence $NE$ has no foreground neighbor anywhere in $N(x)$: upon removal of $x$ it is an isolated foreground pixel, forming its own connected component distinct from any other foreground pixels present.
– *$B_1$, $B_2$ occupy opposite face-neighbors* (e.g. $B_1$ at $N$, $B_2$ at $S$ in Figure B.1b). The remaining two face-neighbors, $E$ and $W$, are not 4-adjacent to each other, and every corner cell is flanked by exactly one of $\{N, S\}$ and one of $\{E, W\}$. Consequently, any foreground cells clustered around $E$ (that is, among $\{E, NE, SE\}$) can only reach foreground cells clustered around $W$ (among $\{W, NW, SW\}$) by passing through $N$ or $S$ — both of which are background. No such path exists, so these two clusters cannot be part of the same 8-connected component.

In both arrangements, the foreground pixels within $N(x)$ split into at least two components that are not 8-adjacent to one another. Building on the connectivity convention established above, write $c(A)$ for the number of connected components of a pixel set $A$ (8-connectivity if $A$ is foreground, 4-connectivity if $A$ is background); we have just shown $c(S \cap N(x)) \geq 2$ in both cases.

Recall that a point is defined to be *simple* exactly when $c(S \cap N(x)) = 1$ (its foreground neighbors form a single connected piece). We have just shown that whenever removing $x$ would merge two background components, $c(S \cap N(x)) \geq 2$ — so $x$ fails the simple-point condition and is never removed. Therefore, no simple-point removal can eliminate a hole.

##### B.5 Equivalence of topology checks

**Theorem 1.** *For 2D Lee94 thinning, the Euler-invariant check is redundant: it is automatically satisfied whenever the border-point and simple-point checks already pass.*

*Proof.* Recall $\chi = O - H$. Consider removing a pixel $x$ that has already passed both the border-point and the simple-point checks:

1. By Lemma B.1 (border-point condition), $\Delta H > 0$ is impossible.
2. By Lemma B.2 (simple-point condition), $\Delta H < 0$ is impossible.
3. Together, these leave only $\Delta H = 0$.

Substituting into the Euler characteristic:

$$\Delta\chi = \Delta O - \Delta H = \Delta O - 0 = \Delta O.$$

The simple-point condition additionally guarantees $\Delta O = 0$ directly (removing a simple point never disconnects the object, by its very definition). Hence $\Delta\chi = 0$ automatically, for every pixel that passes the border-point and simple-point checks — which is precisely what the Euler-invariant check would otherwise be verifying. It therefore adds no further constraint and can be omitted.

##### B.6 Practical implications and considerations

For 2D thinning applications, the theorem above has three direct consequences:

1. No 3D embedding needs to be constructed in an actual implementation — the algorithm can operate directly on a padded 2D image $I \in \{0, 1\}^{(h+2)\times(w+2)}$.
2. Border-point detection simplifies from a 6-neighbor 3D check to the 4-neighbor 2D check: $x$ is a border point iff some $y \in N_4(x)$ is background.
3. The Euler-invariant check can be omitted entirely (Theorem 1); the simple-point check alone suffices for full topology preservation ($\Delta O = \Delta H = 0$), and it can be computed with a direct flood fill over the 8 neighbors of $x$ (collecting the foreground ones and running a depth-first search to count connected components), avoiding the octant/lookup-table machinery the 3D algorithm needs for its Euler-invariant check.
